# Natian and Ryabhatta—graphical user interfaces to create, analyze and visualize single-cell transcriptomic datasets

**DOI:** 10.1101/2021.06.17.448424

**Authors:** Sathiya N. Manivannan, Vidu Garg

**Author notes:** Corresponding Author, Sathiya N Manivannan, Ph.D., Address: 575, Children’s Cross Road, Columbus, OH-43215.

## Abstract

Single-cell transcriptomic analyses permit a high-resolution investigation of biological processes at the individual cell level. Single-cell transcriptomics technologies such as Drop-seq, Smart-seq, MARS-seq, sci-RNA-seq, and CELL-seq produce large volumes of data in the form of sequence reads. In general, the alignment of the reads to genomes and the enumeration of reads mapping to a specific gene results in a gene-count matrix. These gene-count matrix data require robust quality control and statistical analytical pipelines before data mining and interpretation. Among these post-alignment pipelines, the ‘Seurat’ package in ‘R’ is the most popular analytical pipeline for the analysis of single-cell data. This package provides quality control, normalization, principal component analysis, dimensional reduction, clustering, and marker identification among other functions needed to process and mine the single-cell transcriptomic data. While the Seurat package is continuously updated and includes a variety of functionalities, the user is still required to be proficient in the ‘R’ programming language and its data structures to be able to execute the Seurat functions. Hence, there is a demand for a graphical user interface (GUI) that takes in relevant input information and processes the single-cell data using the Seurat pipeline. A GUI will also highly improve the access to single-cell data for life sciences researchers who are not trained in the command-line operation of the ‘R’ platform. To meet this demand, we present R Shiny apps ‘Natian’ and ‘Ryabhatta’ to assist in the generation and analysis of Seurat files from a variety of different sources. The apps and example data can be downloaded from https://singlecelltranscriptomics.org. Natian allows users to create Seurat files from the output of multiple pipelines, integrate existing Seurat files, add metadata information, perform dimensional reduction analysis or upload dimensional reduction data, resume partially processed Seurat files and find cluster markers. Ryabhatta allows users to visualize gene expression using a variety of plotting options, analyze cluster markers, rename clusters, select cells from a graph or based on expression levels of markers, perform differential expression, count the number of cells in each condition, and perform pseudotime analysis using Monocle. We found that the use of these apps substantially improved the analytical and processing time and remove needless troubleshooting due to incompatible commands, typographical errors in scripts, and cluttering of the R environment with variables. We hope the use of these apps improves the use of single-cell data for life sciences research while also providing a tool to learn the functionalities of Seurat and R functions available for single-cell data analysis.

## Introduction

Single-cell/ single nucleus RNA technologies (sc/snRNAseq) allow a detailed and comprehensive analysis of cellular processes in many sub-specialties of life sciences^1–19^. Sc/snRNAseq involves the generation of single-cell or single-nucleus suspensions from tissues or cultured cells and the capture of RNA molecules from individual cells into small compartments. Within these compartments, the cells are lysed, and the RNA molecules are used to generate cDNA with a DNA barcode. The barcode within each compartment is unique and helps in the identification of all cDNA generated from a single compartment^4; 19–22^. This general theme has given rise to a variety of different technologies differing from each other on the method used to create compartments and the target RNA molecules that are captured. Droplet-based techniques such as drop-seq involve the generation of compartments using microfluidics technology with gel beads delivering barcoded oligos and reverse transcription reagents. This technology has been adapted by 10X Genomics to generate the proprietary Chromium controller and capture the 3’ ends of RNA molecules with an oligo-dT mediated reverse transcription^23^. SMART-seq (and the subsequent SMART-seq2) technology can use single-cell capture through fluorescence-activated cell sorting (FACS) or C1-Fluidigm platforms to capture cells in microwell plates and permits the detection of full-length transcripts^19; 24^. Massively parallel single-cell RNA-sequencing (MARS-seq) and Cell Expression by Linear amplification and Sequencing (CELseq) technologies also use FACS to capture individual cells in microwell plates (384-well plates) and provide cost-effective, customizable platforms for detecting RNA molecules^25^. The Seq-well technology employs a portable microwell platform with gel-bead mediated delivery of barcoded oligos to capture single-cells and polyA-tailed transcripts^26^. Single-nucleus sequencing technologies allow for the detection of transcripts from the nucleus of individual cells and are especially useful when the initial sample used is formalin-fixed or paraffin-embedded tissue^22; 27–29^. Single-cell combinatorial indexing (sci-) sequencing technology works with in-situ delivery of barcode labels to fixed cells or isolated nuclei and samples are isolated for second-strand synthesis either using FACS or dilution. The readers are referred to detailed review and benchmarking studies for each of these methodologies elsewhere^12; 30^.

Following the initial barcoding and cDNA preparation, the samples are subject to cDNA library preparation involving ‘sequencing index’ addition in specific single-cell technologies described above and quality assessment of the libraries^31^. The high-throughput sequencing of these libraries results in millions of reads that carry the barcode and the associated RNA sequence information. These reads can be processed using specialized software such as the Cell Ranger pipeline from 10X Genomics^22^ or processed using generic enumerators and alignment tools such as Kallisto, Salmon-Alevin, Bowtie, or STARsolo^32; 33^.

Post-sequencing alignment and counting of reads usually result in a gene versus barcode table with count data for each gene in each cell (the gene-count matrix). Each platform used for alignment and enumeration generates a different format for reporting this data. 10X Genomics Cell Ranger pipeline produces 3 files: the matrix of cell count, the barcode file, and the genes file identifying specific parts of the information. Other methods produce a “dense matrix”, or dgCMatrix format of the gene-count table that can be stored as comma-separated values files (CSV) or tab-delimited text files (.txt) file^32; 33^.

Depending on the technology used, the reads are identified using unique-molecular index (UMI) or read-identity specific to the method^30^. The total of the UMIs or the reads per cell helps identify outliers in the data. These outliers are cells that underwent poor lysis or cells that were partially lysed before the capture of the cDNA, or two or more cells captured within a compartment (doublets) resulting in a substantially higher number of mRNA molecules from a single compartment^2; 21; 30^.

Except in the case of single-nucleus sequencing, RNA molecules coded by nuclear genes as well as cell organelles such as mitochondria and chloroplast can be captured. Genome annotation that includes mitochondrial genes can be used to align and enumerate the genes from both the nucleus and mitochondrial genes. The relative abundance of mitochondrial/other cell organelle genes to that of nuclear-coded genes is usually low in different cell types and therefore can be used as a quality control metric in the evaluation of cellular integrity before sequencing^34^.

The typical output of a single-cell sequencing method ranges from hundreds of cells to thousands of cells. Many single-cell technologies also allow for the sequential and repeated capture of more cells from a single sample source resulting in millions of cells captured and sequenced^30; 35; 36^. With the thousands of genes that can potentially be expressed in each cell type, the gene-count matrix produced from each of these technologies is a voluminous table of millions of data points. These data points represent the gene-count information in a ‘multi-dimensional data plane’ and the data needs to be projected onto a reduced-dimensional space for extracting meaningful information^37^. Before such a dimensional reduction, the counts for each of these cells need to be normalized and scaled across cells in several cases^38–41^. Apart from the count information, many of these experiments require additional information to describe the source of the samples, experimental conditions, and treatments. These are considered as ‘meta’ information and help in evaluating the processed data and extracting information such as differential expression of genes under different conditions.

Several analytical pipelines have been developed for processing the gene-count tables into such manageable and minable datasets. At the time of writing, the most popular (in terms of the number of publications) methods for processing post-alignment data are Seurat and Monocle^39; 42^. Seurat is an R package that enables users to perform quality control, normalization, dimensionality reduction, clustering of cells among several other functionalities^38; 39; 43; 44^. These functionalities are written into facile and versatile functions that are easy to learn and implement for bioinformaticians and computational biologists across life sciences disciplines. Monocle is an R package that has also been used in a large number of studies to process and analyze single-cell data. Additionally, Monocle provides an additional tool to analyze the temporal changes in samples using the pseudotime analysis step^45–47^. The details of each of these steps taken to perform normalization, scaling data, principal component analysis, dimensional reduction, and pseudotime analyses are well documented through several publications and dedicated web-portals^48; 49^.

However, with a large number of life sciences researchers, the analysis of single-cell sequencing data has been limited due to the command-line nature of these tools. While functions and steps used for Seurat and Monocle are easy to learn for researchers with an R background, the functions and data formats are complex enough to be a barrier for the exploration of data by non-computational scientists. Learning of R and the tools associated with single-cell data analysis is the ultimate solution to the researchers’ interested long-term analysis of the scRNAseq data, but the exploration of the use and scope of single-cell datasets should not be limited by computer literacy. To bring a computational and non-computational scientist to a common table, to be able to analyze the data and discuss the result of their analysis, we provide two graphical user interphases (GUIs) ‘Ryabhatta’ and ‘Natian’. These GUIs are based on the Shiny package in R and are provided to users with no R background. The installation and running of these GUIs have been evaluated in Windows, macOS, and Linux operating systems. The inclusion of life sciences researchers with extensive experience in molecular and cellular biology, but limited R experience in single-cell data analysis will provide novel biological insights on existing and future single-cell data sets.

## Results

### Standard processing of a single Single-cell dataset using Natian

We developed Natian to streamline the processing of single-cell data from the post-alignment output to the processed Seurat files without the need for command-line inputs (Figure 1). The shiny application provides the opportunity (or ability) to get data from 10xGenomic output which is present in the ‘outs/filtered_gene_bc_matrices’ folder. Alternatively, users can also use the Gene Expression Omnibus submission IDs (‘GSE…’) to identify the processed count-matrix files using the GEO button. The data can be uploaded in a compressed format such as ‘gzip’ and the count matrix can be one of the following formats: CSV or txt. Alternatively, an ‘RData’ file with the dGEcount matrix can also be uploaded. Upon successful upload of either of these kinds of data, Natian provides an option to name the Seurat file. Currently, only alphanumeric values (A-Z, a-z, 0-9) characters with no space or special characters are permitted in the name of the Seurat file. An exception is included for an underscore (“_”). The name may not start with a number. These restrictions are due to the limitations in the variable names in R statistical platform. A Seurat file is now created in the R environment. Next, an option to add metadata is presented which can be used to enter the treatment, time point/developmental stage, gender, age, sample, replicate, ID, or other information relevant to the Single-cell data. In some cases, the single-cell data submitted or processed may have all the relevant information stored as column names in the dataset. In this case, generated metadata using the column attribute can be used to get the meta-information.

**Figure 1.**
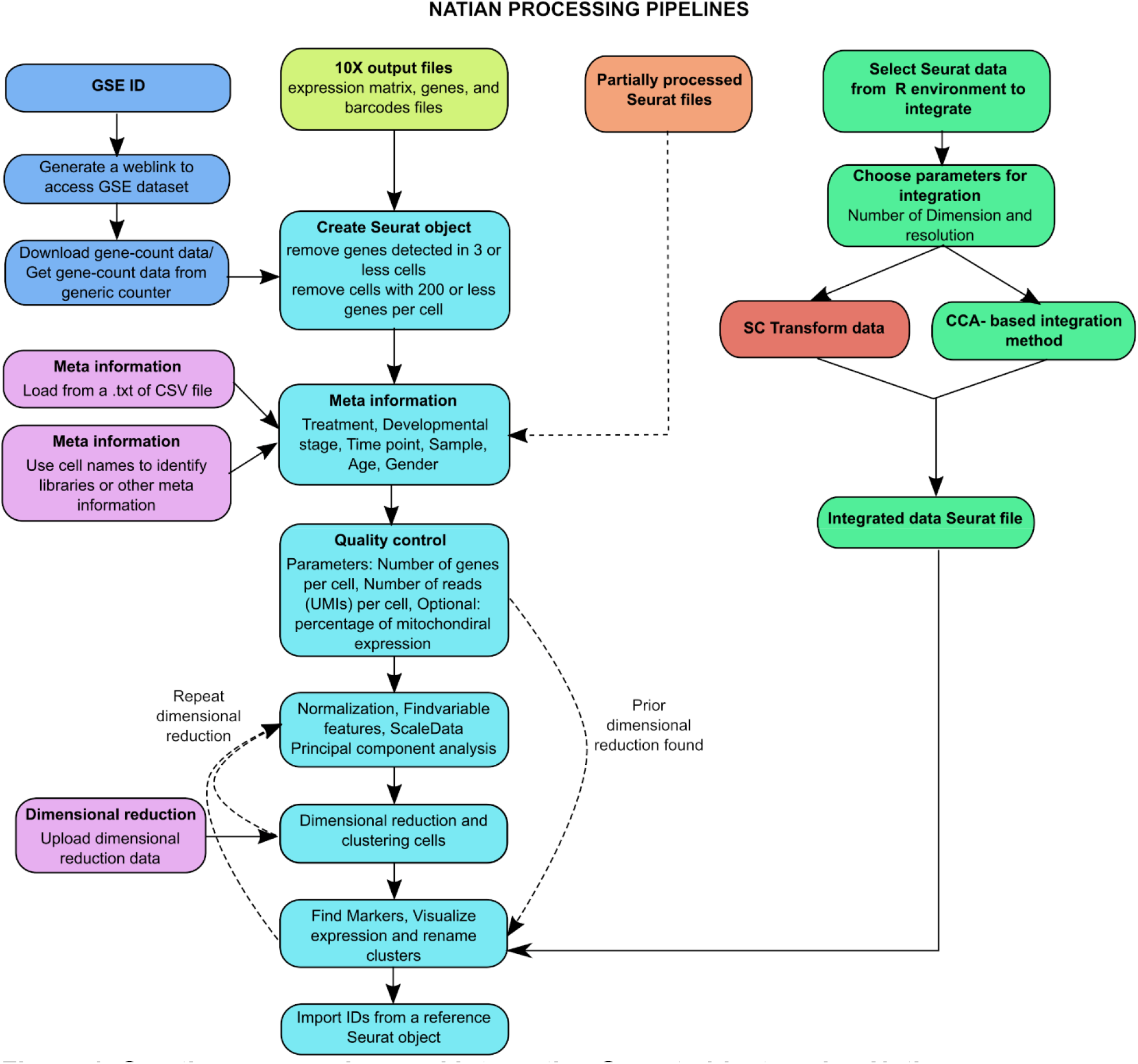
Creating, processing, and integration Seurat objects using Natian. Flow diagram showing the sequence of steps involved in creating a Seurat object from the single-cell output from 10x Genomics or other pipelines and integration of existing Seurat objects. Element/steps are color-coded to show common (cyan), GEO data analysis (blue), 10x output analysis (lime green), partially processed (orange), integration of Seurat object (green), optional (purple), and scTranformation step (red). Solid lines indicate the normal flow of steps, dotted lines indicate alternative routes.

The next step when performing the analysis using the standard pipeline is to measure quality control metrics. The number of genes and the number of reads captured in each cell form the standard metrics to identify outliers. Outliers with high levels of either of these two parameters are likely doublet cells and need to be removed. Cells with low levels of either of the parameters likely suffered from poor capture or sequencing artifacts. In some cases, the use of the percentage of mitochondrial transcripts registered compared to all the reads within a cell also assists in the identification of cells with a higher level of background or partial capture of nuclear transcripts. Natian provides options to iteratively check the limits of each of these parameters to determine the optimal thresholds with feedback on the number of cells filtered through the process. Once this step is done, Natian proceeds to normalize the data, find variable features, scale the data and perform principal component analysis. The result of these processes is an ‘elbow’ plot that shows the changes in the standard deviation versus the number of dimensions (‘dims’) used. This heuristic approach allows users to determine the number of dims to use for dimensional reduction, finding neighboring cells in the graph-based clustering analysis. The second graph presented below the elbow plot is a Clustree diagram, which displays the number of possible clusters obtained by clustering the data using the Louvain algorithm with a range of resolution parameters in Seurat. Once a suitable number of dims and resolution is determined, the dimensional reduction process creates a Uniform Manifold Approximation and Projection (UMAP) plot with the clusters determined according to the resolution provided. Simultaneously, a cell count chart listing the number of cells in each cluster is created. The user has the option to repeat the clustering and dimensional reduction with a different number of dims and resolution parameters until the desired number of clusters is obtained.

The markers that are significantly enriched in a specific cluster can be determined using the find markers options. Natian uses the default Wilcox test to perform this analysis using thresholds for differential expression change (logfc.threshold) and the minimum fraction of cells (min.pct) that need to be positive for the gene expression. The output shows top markers in a table format and using the switch to markers view, the marker expression can be visualized in the UMAP plot on the top.

### Alternative routes towards getting metadata and dimensional reduction

Metadata can be added from a CSV or txt file format. Such data can be imported instead of manually entering a value for each category. This feature is compatible if an initial analysis was done using a different tool such as the loupe browser from 10X Genomics. After the addition of metadata dimensional reduction, the coordinates from a previous run/publication can also be imported to the Seurat object. This might be convenient for users who do not wish to repeat the analysis or would like to follow the dimensional reduction analysis performed by the authors/investigators of the original publication or run. This step also performs the cell removal according to the available dimensional reduction data.

### Integration of single-cell datasets and resuming processing of Seurat files using Natian

Integration of single-cell datasets is essential to perform the analysis of multiple runs of single-cell data or data derived from different conditions. Batch correction is also an additional step performed to remove differences between single-cell datasets owing to the depth of sequencing or variations in capture. To facilitate this integration and batch correction, users can use the ‘integrate Seurat’ button in Natian; and select the files to integrate, input the number of dimensions to use, and optionally choose sc-transformation of individual samples before integration^41^. A standard dimensional reduction pipeline and clustering are performed on the integrated data. The integrated data can be subject to refined clustering and dimensional reduction using the resume Seurat processing pipeline.

The ‘Resume processing Seurat file’ button is useful to resume processing of partially processed Seurat files. It can also be used if the Seurat object was created in a previous version or if the addition of metadata, external dimensional reduction analysis, or a rerun of clustering is required. The data analysis pipeline determines if different steps (principal component analysis, dimensional reduction, etc.,) have been performed and directs the user to the incomplete steps in the pipeline described in the standard processing section.

### Loading processed Seurat files in Ryabhatta and initial options

Once a Seurat dataset is processed using Natian or a conventional R script/command line process, it can be used to visualized using Ryabhatta (Figure 2). Ryabhatta opens with an option to load an “.RData” or “.RDS” format files with the processed Seurat files into the R environment. Processed Seurat files in the R environment can be loaded using the load ‘Seurat object’ button. Once processed Seurat files are loaded, a central dimensional reduction plot is displayed along with cell number count, and other processing/visualization options shown in the various panels.

**Figure 2.**
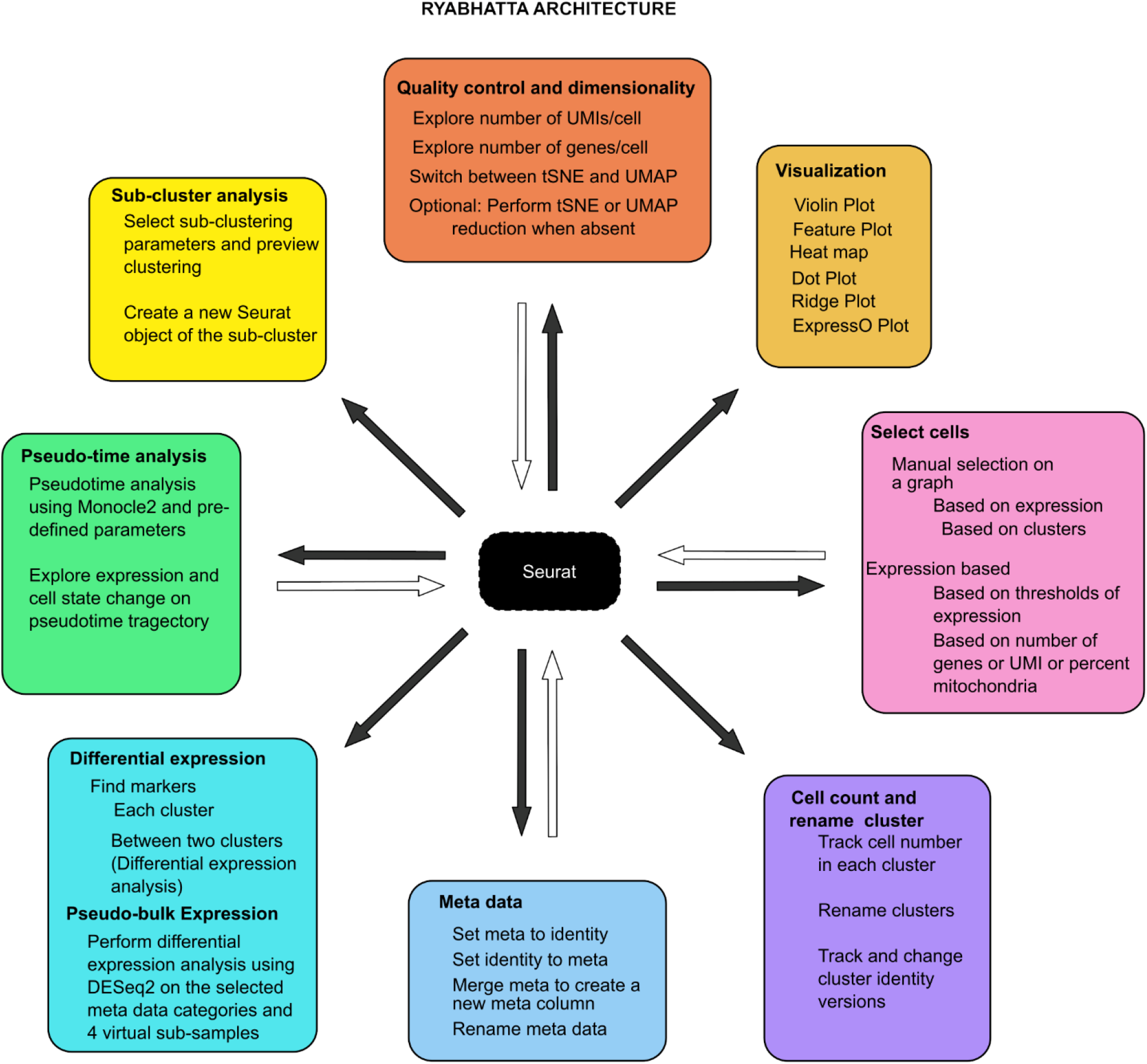
Analysis, exploration, and visualization of Seurat using Ryabhatta. Network diagram showing the various visualization and analysis steps possible using Ryabhatta. The analysis uses a central Seurat object that is mined for different data analysis steps. Steps that extract information from the Seurat object are shown connected to the object with a black arrow. Steps that return information to the Seurat object or modify the Seurat object are shown connected through a white arrow.

Ryabhatta sets the default analysis to the “RNA” assay of the Seurat object. The genes present in the ‘scaledata’ slot are used for visualization. If scaled data is not available for the “RNA” data slot, data are scaled before the options are loaded. If UMAP or tSNE dimensional reduction is performed, an error message is displayed, and the users have the option to perform dimensionality reduction using UMAP or tSNE algorithms with Ryabhatta. It is preferable to use Natian if such an error is encountered to make sure the data set is processed in a way suitable for analysis with Ryabhatta.

On top of the dimensional reduction plot, the name of the current Seurat file is displayed with summary information as a tooltip. Below this, a set of buttons provide for the use of different visualization options. This includes the number of UMIs, the number of genes per cell, and if different classes of metadata are available, these data can also be visualized. The metadata dropdown menu on the left panel allows the user to select metadata such as the developmental stage to control the separation of data plots and clusters. An option to change existing metadata as cluster identities or vice versa is available from the buttons above the dimensional reduction plot.

It is possible to change the visualization option from UMAP to tSNE and back using these buttons. If a tSNE dimensional reduction has not been performed, this could also be done within Ryabhatta by selecting the perform tSNE reduction with appropriate dims. The ‘download plot’ button lets the user download a high-resolution PDF of the current plot in this panel. The cell counter graph on the right panel displays the number of cells in each of the current identities. This data can be downloaded as a CSV file using the download cell count button.

### Visualization options in Ryabhatta

Expression in the “RNA” slot of the data can be visualized using the variety of plots available in Ryabhatta. This includes a violin plot with an option to show the individual points; a feature plot showing the expression overlaid on the chosen dimensional reduction (tSNE/UMAP); a heatmap to show the expression of all the cells in the form of a heatmap; dot plot where expression and percentage of cells expressing a specific gene can be visualized; ridge plot to show expression as stacked ridges; and an expresso plot which permits multiplex expression analysis of up to 3 genes.

In each case, the plot may be split according to metadata in the metadata option. The default Seurat functions for the following plots have been modified: the violin plot has been altered to permit the display of points in the violin according to the color of the cluster/condition and shows mean, and 5^th^ and 95^th^ quantiles in the plot. The dot plot has been modified to show a color guide scale and displays expression for each gene. Any plot drawn using these options can be downloaded using the download plot option in the Plot options panel. The colors of the feature plot or the violin plot can be edited before downloading by using the settings tab next to the plot area.

### Selecting cells in Ryabhatta

Cells can be selected using one of the two selection options. 1) A graphical selection method that allows users to draw boundaries using a mouse cursor around cells of interest. In this method, the users can select a gene’s expression as their guide or select no genes to get a dimensional reduction plot with the current identities labeled in the plot. Manual selection can be followed by renaming the selected cells with a unique identity. 2) An expression-based selection method that allows users to select cells based on expression thresholds set for various genes in the ‘RNA’ slot or the total number of genes per cell (‘nFeature_RNA’) or the total number of UMIs per cell (‘nCount_RNA’) or the percentage of mitochondrial transcripts among total reads (‘percent.mito’) options. The number of cells selected in the expression-based option will be displayed along with the dimensional reduction plot, highlighting selected cells in red in the plot area of Ryabhatta. Additionally, a specific cluster can be chosen to be the focus of this selection. Users can use this option to select cells that express a specific set of markers within a chosen cluster. Cells thus chosen can be given a new name or used to create a new Seurat object.

### Differential expression analysis in Ryabhatta

Ryabhatta provides two options to perform differential expression analysis. 1) The canonical ‘findmarkers’ function in Seurat-based analysis. This comes with the options to choose different test methods including ‘wilcox’, ‘bimod’, ‘t’, ‘roc’, ‘poisson’, ‘negbinom’, ‘DESeq2’, ‘MAST’, and ‘LR’.

Choosing 1 cluster or no clusters will result in the identification of markers for all the clusters using the thresholds set in this panel. After the completion, the top 5 markers of each cluster are shown in the form of a heatmap. All identified markers in individual clusters are shown at the bottom in a collapsible table. Users can use the ‘switch to markers’ button view to evaluate the expression of each of these markers overlaid on the dimensional reduction plot. Users can also download the marker data as a ‘.csv’ file which includes the following columns:

p_val: P-value obtained from the analysis using the chosen test.
avg_logFC: natural log-transformed fold change in expression between a given cluster compared to all the other clusters combined.
pct.1: Percentage of cells with detectable levels of expression within the cluster
pct.2: Percentage of cells with detectable levels of expression outside the cluster
p_val_adj: adjusted P-value
cluster: the identity of the cluster.

If two specific clusters are chosen, these will be used to calculate differential expression between the chosen clusters. The output is a volcano plot of -log10(p_val) vs avg_logFC of the gene expression and the size of the points as a function of the ratio of pct.1/pct.2.

2) A pseudobulk differential expression analysis is also provided in Ryabhatta which is based on splitting the cells within a cluster into 4 virtual samples and combining the counts for the expression of each gene within individual virtual samples. This approach mimics bulk RNA-seq analysis data collection and therefore can compensate for high levels of dropout observed in single-cell data technologies. We have observed that the number of genes identified using pseudobulk technology is significantly higher than the ‘Wilcox’ test method using the FindMarkers. This includes genes that might have been removed due to the percentage expression threshold. The pseudobulk differential expression analysis works using the metadata categories as conditions and clusters as individual samples and uses DESeq2 to obtain identify differentially expressed genes. The normalization step in the DESeq2 pipeline is used to adjust expressions for differences in cell number between conditions.

The output of this analysis is also a volcano plot of log2FoldChange vs -log10(pvalue) and a .csv file. The CSV file contains the standard DESeq2 output values such as ‘baseMean’, ‘pvalue’, ‘padj’, and ‘log2FoldChange’ along with normalized counts per million reads of each gene in each virtual sample. This will permit the comparison of single-cell RNA seq data to bulk RNA-seq data and cross-platform analyses.

### Subclustering analysis using Ryabhatta

Ryabhatta has sub-cluster analysis built into it. By selecting the cluster(s) users can focus on specific cell types within the data for further analysis. Sub-clustering analysis subsets specific data sets and uses an initial estimated number of dimensions to create temporary dimensional reduction and clustree diagram showing changes in the number of clusters within the subcluster at different resolutions. A key feature of this sub-clustering analysis is that it will use an integrated assay to perform this analysis if the data had been created using Seurat integration. It also removes low expression genes (less than 3 cells with detectable expression) and cells with low gene expression (with less than 200 genes/cell) from the selected cluster. With the input of desired resolution and dimensions, a new sub-cluster is created bearing the name of the original Seurat file and clusters used to create the sub-cluster. This method of sub-clustering is also used when manually selected cells or cells selected based on expression are chosen for sub-clustering analysis. The rename Seurat button at the start of the Ryabhatta app can be used to change the name of any of the Seurat objects.

### Monocle 2 Pseudotime analysis using Ryabhatta

Ryabhatta uses Monocle 2 to perform pseudotime analysis by using the raw counts of the cells in the RNA assay slot of the Seurat object. Then the data is filtered to remove genes with a minimum expression cutoff of 0.1 and genes that show expression in less than 50 cells. A standard analysis pipeline is then employed to obtain the dimensional reduction and graph parameters using the DDRTree approach. These data are returned to the Seurat object and stored as additional features.

Once this analysis is complete, the users can use Ryabhatta to visualize gene expression, Monocle BEAM-based pseudotime ‘states’, and calculated pseudotime using the ‘Plot pseudotime’ option. The information from the Monocle2 analysis is stored as metadata in the Seurat object. This allows the comparison of pseudotime states with that of the Seurat clustering analysis results.

### Convenience tools in Natian and Ryabhatta

Natian provides an inbuilt Note-taker tool to actively register pertinent information about the processing of the Seurat file using a note-taker tool. This tool continuously monitors and adds processing steps and information about cells filtered or data upload files used in the processing of the single-cell data. A ‘download’ button enables collecting the session notes in a facile HTML file with the graphs and other data embedded in a single file. This information will be valuable for post-process evaluation or in generating a process report.

Ryabhatta allows users to import genes from a text file or CSV file to analyze them together. Gene lists used in Ryabhatta can also be saved and recalled using the load genes panel. This is especially useful to generate a heatmap of many genes or analyze a set of genes in different Seurat objects. Genes are filtered to make sure only expressed genes are used for the analysis. Ryabhatta also allows users to combine metadata fields to generate a new metadata field. This allows on-the-fly examination of different combinations of conditions such as developmental stage, timepoint, or treatment methods. Ryabhatta also has the option to seamlessly transfer the metadata information to the identities of the cells and add identities to the metadata information.

Ryabhatta allows users to save the previous version of identities. With every rename or change of cluster identity, a new identity version ‘Identversion’ is created that is stored with the Seurat object. This can be used to navigate between differently named versions or revert to an earlier version of cell identities.

### Requirements and testing

Ryabhatta and Natian have been tested on desktops and laptops with 4 GB to 32 GB memory in Windows and Macintosh operating systems. We have also tested the use of Ryabhatta and Natian on Amazon cloud computational services including EC2 and T2 tiers. The apps are currently limited to 20 GB of individual file size. This corresponds to ~80,000 cells with ‘scaled.data’ for ~19,000 genes. In our experience, a vast majority of the analysis that is performed on personal laptops and desktops rarely exceeds 40,000 cells. With the implementation of Ryabhatta and Natian on high computational devices, it may be possible to remove this limitation.

Rybhatta and Natian have been developed on Seurat 3.2.1, R version 3.6, and RStudio 1.0. Newer versions of R and RStudio have been used to evaluate the performance. We have also tested Seurat version 4.0 with these apps. Users will have to install R and RStudio on their devices. Windows users will need Rtools (Rtools 30 or newer) to use Monocle and associated functions. Macintosh users may have to install an XML package from a binary and authorize this installation. In Windows, apart from R and Rstudio, users also need to install Rtools. This is also available from r-project.org. In Macintosh OSX machines, the users will need to install Xquartz to enable the use of the Cairo package for downloading high-resolution PDF files. The new version of OSX also disables the Arial font. This needs to be enabled using the Fontbook application on Macintosh machines.

### Source code, test data, and user manual

Proof-of-concept analysis using the apps for integrating multiple data sets, performing subcluster analysis as well as differential gene expression analysis can are provided in the supplementary data. (Supplementary Files 1, 2, Supplementary Figures 1,2, Supplementary Tables 1, 2).

Video tutorial and test data are available online at https://singlecelltranscriptomics.org.

## Discussion

The GUIs we present here are not be intended to replace a bioinformatician or curb the enthusiasm to learn command lines but to remove the requirement to learn command-line to explore single-cell data. Life science researchers without R background can now explore the single-cell data generated and processed into Seurat files. The apps also provide a window into the steps involved in the processing of single-cell data to non-computational biologists and the iterative exploration of different parameters to process the data. We believe this will eventually lead to an interest in learning command-line programs to further expand the analytical tools provided in the GUIs.

Several publicly available browser-based tools such as the UCSC cell browser, cellxgene, and Broad Institute single-cell portal allow for analysis of data that has been loaded into respective portals. However, users cannot perform sub-cluster analysis or provide visualization methods that are available with Ryabhatta. Commercial tools, such as the 10x Genomics’ Loupe browser and Bioturing, offer limited ability to modify processing pipelines, subclustering analysis, and integrating datasets in Seurat format. Similar R Shiny-based apps such as SCHNAAP, CIPR do not work on preserving the Seurat object. Others such as Seurat Wizard do not permit sub-cluster analysis or provide additional visualization tools provided in the Ryabhatta. Both Ryabhatta and Natian preserve the Seurat object and all analyses are stored in the Seurat object including the Monocle-based pseudotime analysis. This allows for saving and sharing of the processed Seurat object for subsequent analysis or pipeline development.

From the inception of the development of these apps, we made a conscious decision to make sure that every element of the program is accessible to the user for edits and modification. Therefore, all the functions, java-script, jQuery, and HTML commands that have been used in the apps are accessible in the shiny app. This allows users to tweak steps that might be specific to their needs or co-opt scripts and functions in the app to develop additional pipelines.

Finally, these apps are possible due to the continued development of Seurat and Monocle and other additional R packages developed by a large community of dedicated bioinformatics researchers. We believe that these GUIs shine a light on the importance of these tools and the requirement for further development and expansion of such tools.

## Acknowledgments

We would like to thank Dr. Madhumita Basu, Dr. Dwitiya Sawant, Dr. Brenda Lily and Dr. Uddalak Majumdar at the Center for Cardiovascular and Pulmonary Research and Summer Fair, Dr. Tracy Bedrosian, James Fitch at the Institute for Genomic Medicine in Abigail Wexner Research Institute at Nationwide Children’s Hospital for testing the Shiny apps as well as providing valuable feedback. We are grateful for the technical support by Mr. Sabari Ayyappan Manivannan and Mr. Sivaganesh Manivannan for their help in designing and publishing the webpage for the distribution of the source code and video manual. S.M was supported by a T32HL098039-06A1 training award from the National Institutes of Health, a postdoctoral fellowship award from the American Heart Association and the Children’s heart foundation (Award number 821190), and the postdoctoral idea award from the Abigail Wexner Research Institute at the Nationwide Children’s Hospital (AWD00001356).

## Notes

https://singlecelltranscriptomics.org

### Competing Interest Statement

The authors have declared no competing interest.

