## Supplementary figures and images for "Natian and Ryabhatta—graphical user interfaces to create, analyze and visualize single-cell transcriptomic datasets"

### Supplementary Figures 1-2

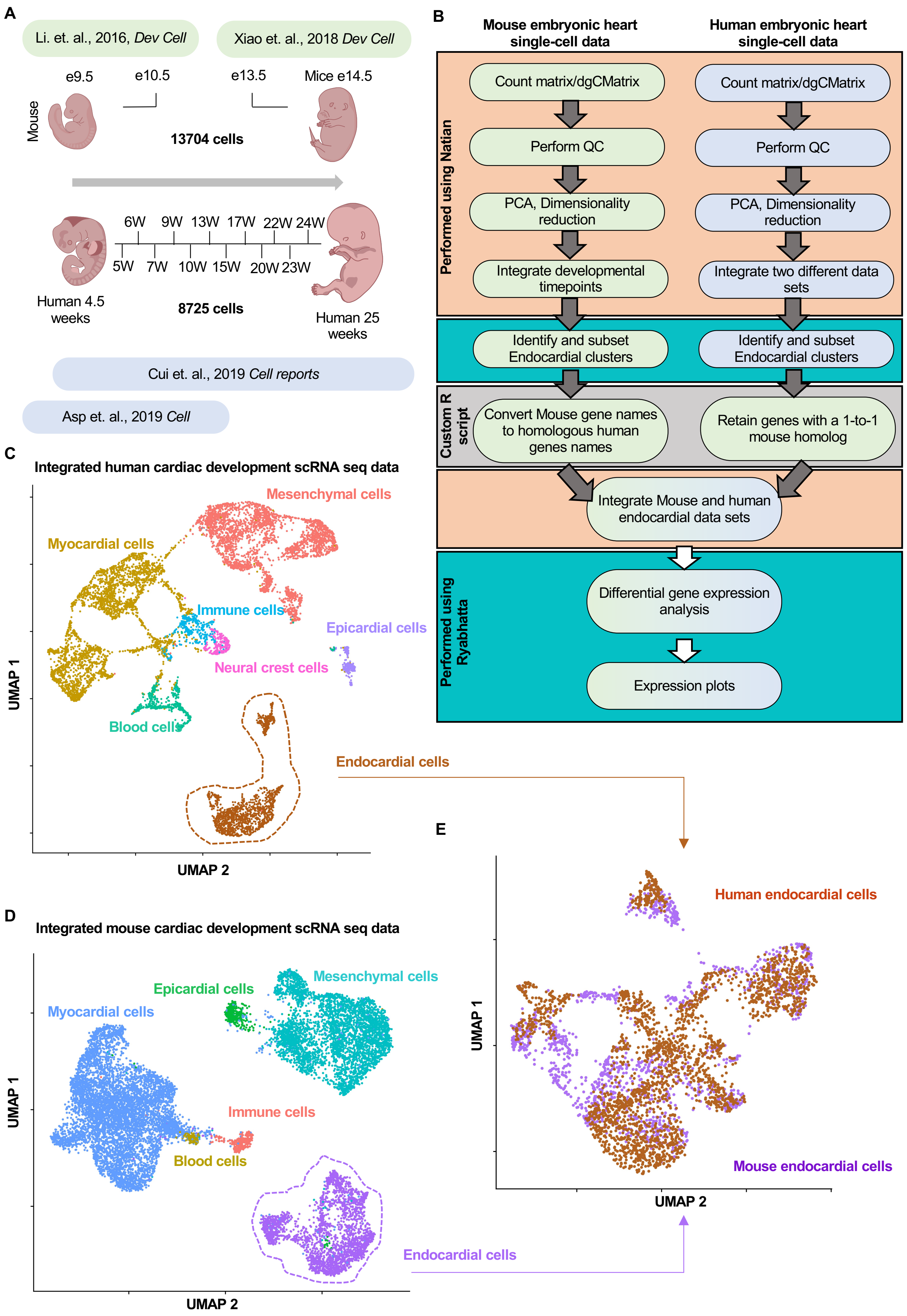

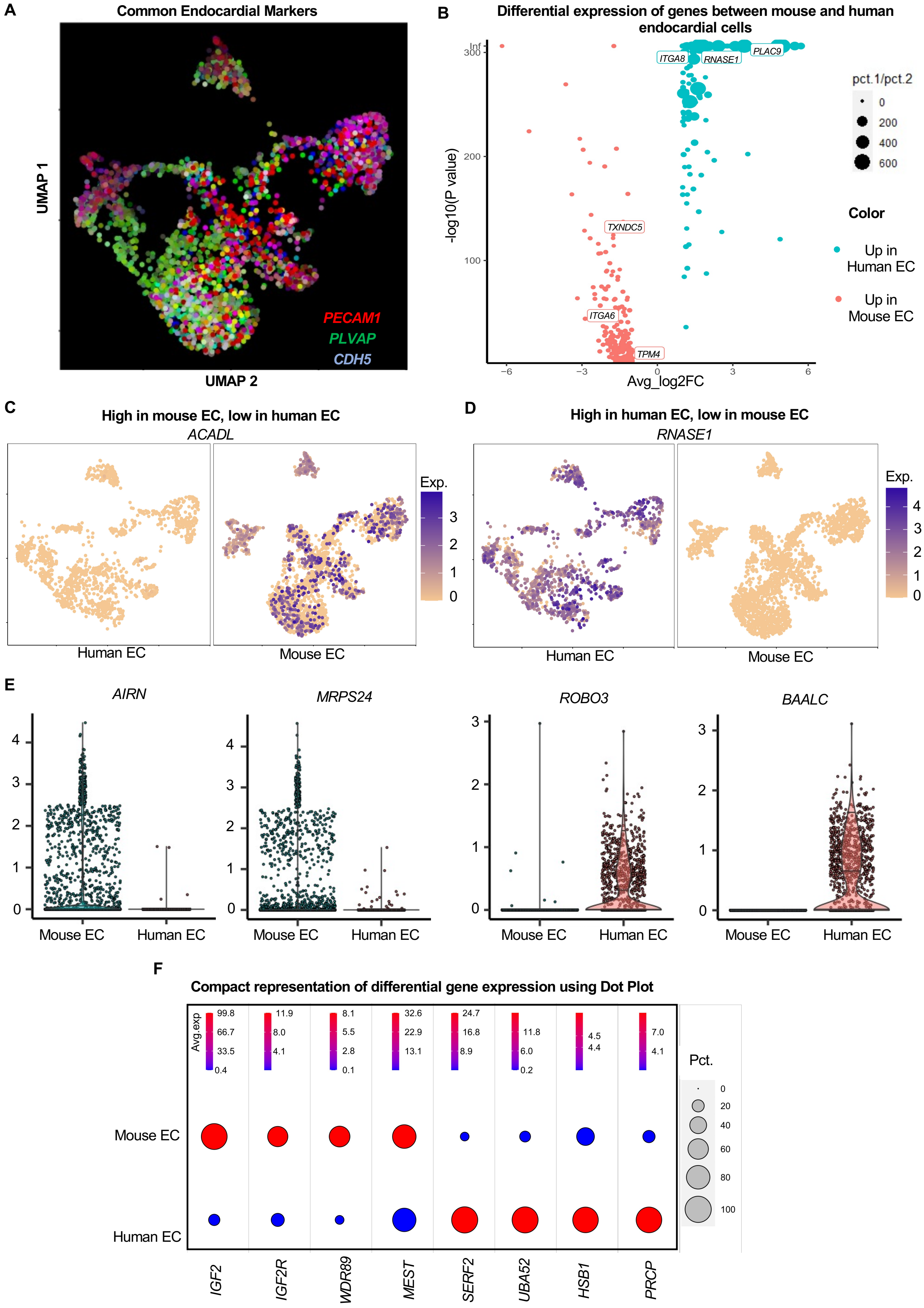
