## Supplementary File 2 for "Natian and Ryabhatta—graphical user interfaces to create, analyze and visualize single-cell transcriptomic datasets": R script mouse to human gene names.R

```
#Author: Sathiya N. Manivannan
#Goal: To convert the gene names of a seurat object from mouse gene names to homologus human gene names
#Note: This method only evaluates one to one homologs between a mouse gene and a human gene name.
#Note: Seurat objects are reduced to only genes where one-to-one homologs are available, rest of the genes will cause
#false positive during differential expression due to the lack of homologus counterparts.
Change_names <- function(Mouse_Seurat, Human_Seurat){
  library(Seurat)
  HMD_HumanPhenotype <- HMD_HumanPhenotype[HMD_HumanPhenotype$Human_gene_symbol %in% row.names(Human_Seurat),]
  Human_Seurat <- subset(Human_Seurat, features = HMD_HumanPhenotype$Human_gene_symbol)
  Mouse_Seurat <- subset(Mouse_Seurat, features = row.names(HMD_HumanPhenotype))
  reductions <- Mouse_Seurat@reductions$umap@cell.embeddings
  meta_data <- Mouse_Seurat@meta.data
  misc <- Mouse_Seurat@misc
  idents <- Idents(Mouse_Seurat)
  counts = as.matrix(Mouse_Seurat@assays$RNA@counts)
  row.names(counts) <- HMD_HumanPhenotype[row.names(counts),]$Human_gene_symbol
  new_Seurat <- CreateSeuratObject(count = counts, min.cells = 0, min.features = 0)
  new_Seurat <- NormalizeData(new_Seurat)
  new_Seurat <- FindVariableFeatures(new_Seurat)
  new_Seurat <- ScaleData(new_Seurat)
  new_Seurat <- RunPCA(new_Seurat)
  new_Seurat[["umap"]] <- CreateDimReducObject(reductions, key = "UMAP", assay = "RNA")
  Idents(new_Seurat) <- idents
  new_Seurat@misc <- misc
  new_Seurat@meta.data <- meta_data
  assign("Mouse_EC_homolog",new_Seurat, envir = .GlobalEnv)
  assign("Human_EC_homolog", Human_Seurat, envir = .GlobalEnv)
  return(NULL)
}

Volcanoplot <- function(DEG_output, label_genes){
library(ggplot2)
library(ggrepel)
library(dplyr)
#filtereing genes based on adjusted P value and avg_log2FC threshold
DEG_output <- DEG_output[DEG_output$p_val_adj <= 0.05 & abs(DEG_output$avg_log2FC) > 1,]
row.names(DEG_output) <- DEG_output$X
DEG_output <- DEG_output[complete.cases(DEG_output),]
#calculating -log10 transformed P values
DEG_output$log10Pvalue <- -log10(DEG_output$p_val_adj)
#adding up/down labels to genes depending on the average log2 fold change
DEG_output$up_down <- ifelse(sign(DEG_output$avg_log2FC) > 0, "Up", "Down")
#getting maximum finite values
max_val <- max(DEG_output$log10Pvalue[is.finite(DEG_output$log10Pvalue)])
#fixing infnity values created during -log10 transformation of P values for Volcano plots.
DEG_output$log10Pvalue[!is.finite(DEG_output$log10Pvalue)] <- sign(DEG_output$log10Pvalue[!is.finite(DEG_output$log10Pvalue)])*(abs(max(DEG_output$log10Pvalue[is.finite(DEG_output$log10Pvalue)]))+5)
#Creating a labelling dataframe
DEG_label_df <- DEG_output[label_genes,]
DEG_label_df$labels <- row.names(DEG_label_df)
#plotting genes
gg <- ggplot(DEG_output, aes(y = log10Pvalue, x= avg_log2FC, color = as.factor(up_down), size = pct.1/pct.2))+geom_point()+
               geom_label_repel(data = DEG_label_df, aes(y = log10Pvalue, x= avg_log2FC, color = as.factor(up_down), label = labels),size= 2, inherit.aes = F)+
               scale_y_continuous(breaks = unique(c(seq(from= 0, to = max_val, by= 100), max_val+5)),
                                  labels = as.character(c(seq(from= 0, to = max_val, by= 100), Inf)),
                                  expand = expansion(mult = c(0, 0.02)))+
  theme(axis.line = element_line(size= 0.8),
        axis.title = element_blank(),
        axis.text = element_text(size= 8),
        panel.background = element_blank(),
        plot.background = element_blank(),
        panel.grid = element_blank())
return(gg)
}
```
